# Chlamydial protease-like activating factor targets SLC7A11 for degradation to induce ferroptosis and facilitate dissemination

**DOI:** 10.1101/2023.12.20.572489

**Authors:** Wentao Chen, Xin Su, Yuying Pan, Yaohua Xue, Lihong Zeng, Qingqing Xu, Xueying Yu, Xiaona Yin, Han Zhou, Zhanqin Feng, Bao Zhang, Wei Zhao, Heping Zheng

## Abstract

*Chlamydia trachomatis*, the most prevalent bacterial agent of sexually transmitted infections, possesses remarkable capacities for dissemination within the host, leading to reproductive health complications. The release of progeny through the orchestrated lysis of host cells plays a crucial role in *Chlamydia* dissemination, but the underlying molecular mechanisms remain largely elusive. Here, we uncovered a novel mechanism by which *Chlamydia* induces host cells ferroptosis to facilitate its dissemination. This process involves the degradation of host protein SLC7A11 by the chlamydial protease-like activating factor (CPAF), resulting in glutathione depletion, oxidative damage, and subsequent host cell lysis characterized by lipid peroxidation. Infection with CPAF-deficient strain fails to induce host cells ferroptosis, leading to restricted progeny release. Importantly, inhibiting ferroptosis effectively limits the release of *Chlamydia* progeny, highlighting its potential as a therapeutic strategy for controlling *Chlamydia* dissemination. These findings provide insights into the chlamydial conserved dissemination strategy and enhance understanding of its pathogenesis.

## Introduction

*Chlamydia trachomatis* (CT), an obligate intracellular pathogen, is responsible for the most common bacterial sexually transmitted diseases. Untreated *Chlamydia* infections can result in severe reproductive health complications, including pelvic inflammatory disease (PID), ectopic pregnancy, infertility in women (1, 2). These complications associated with *Chlamydia* is inseparable to pathogen’s successful dissemination and propagation within the body.

CT exhibits a distinctive biphasic life cycle involving the invasion of host cells by infectious elementary bodies (EBs), their transformation into replicative reticulate bodies (RBs) within specialized intracellular compartments called inclusions, and subsequent re-differentiation back into EBs (3). The release of progeny through host cell lysis and their subsequent initiation of new infections is a critical process for the dissemination and propagation of CT (3, 4), implying that blocking this process could be an effective intervention to disrupt the spread of infection. However, the underlying molecular mechanism of host cells lysis driving by CT, are still elusive.

Cell death modalities observed during pathogen infection play a dual role in the intricate interplay between pathogens and hosts. They can either serve as beneficial outcomes for host defense against invading pathogens or be exploited by pathogens to enhance their pathogenicity and promote dissemination (5). Ferroptosis (6), a newly characterized form of programmed necrotic cell death resulting from impaired lipid peroxide repair systems, exerts significant influence on diverse physiological processes and various disease conditions, including infectious diseases (7-12). Although the involvement of ferroptosis in the pathogenesis of CT has not been demonstrated so far, the replication of *Chlamydia* leads to the generation of reactive oxygen species (ROS), which subsequently causes membrane lipid peroxidation (13). Moreover, supplementing *Chlamydia*-infected lambs with vitamin E, a potent inhibitor of ferroptosis, results in improved treatment outcomes (14). The potential interplay between CT and host cells in relation to ferroptosis remains an area requiring further investigation, as suggestive evidence exists but the specific mechanisms and involvement of ferroptosis in *Chlamydia* infection have not been fully elucidated.

Chlamydial protease-like activity factor (CPAF) is a secreted protease produced by CT and other *Chlamydia* species. Functioning as a bacterial virulence factor, CPAF exhibits the ability to cleave host cell proteins, including cytoskeletal proteins (15), transcription factors (16), and immune signaling molecules (17). This proteolytic activity allows CPAF to undermine host defense mechanisms, facilitate chlamydial survival and replication, and evade host immune surveillance (15-19). While CPAF has been implicated in the cell lysis process (18, 20, 21), the specific mechanism by which it operates remains poorly understood. In this study, we uncovered the critical role of CPAF in facilitating microbial dissemination by inducing host cell ferroptosis through the degradation of SLC7A11. This process led to a depletion of glutathione, accumulation of lipid peroxidation, subsequent lysis of host cells, and release of progeny. Deficiency of CPAF or pharmacological inhibition of ferroptosis proved to be effective in limiting the dissemination of CT. These findings provide valuable insights into the pathogenesis of *Chlamydia* and its conserved dissemination strategy.

## Results

### *Chlamydia* induces ferroptosis in host cells to facilitate the release of progeny

To investigate the lysis process during infection, we employed a model using HeLa-229 cells infected with CT serovar D (CT-D) at different multiplicities of infection (MOIs). We performed a time-course analysis to evaluate cell death by utilizing propidium iodide (PI) staining and lactate dehydrogenase (LDH) release assays. The results demonstrated that CT caused substantial cell death during the late stages of infection (Fig. 1a and b). Accumulation of lipid peroxides is a characteristic feature of ferroptosis (6). To explore the induction of lipid peroxide accumulation by CT during the late stage of infection, we quantified the levels of lipid peroxidation in CT-infected cells using C11 BODIPY 581/591. Our findings revealed a significant time- and MOI-dependent increase in lipid peroxide levels within the infected cells (Fig. 1c). Next, we examined whether the CT-driven cell death could be rescued by ferroptosis inhibitors, specifically ferrostatin-1 and liproxstatin-1, known to prevent lipid peroxide-mediated ferroptosis (22). Treatment with either compound reduced the numbers of PI-positive cell, prevented LDH release, and attenuated lipid peroxide accumulation (Fig. 1d and e), indicating that CT induces ferroptosis during its late developmental cycle.

**Fig. 1.**
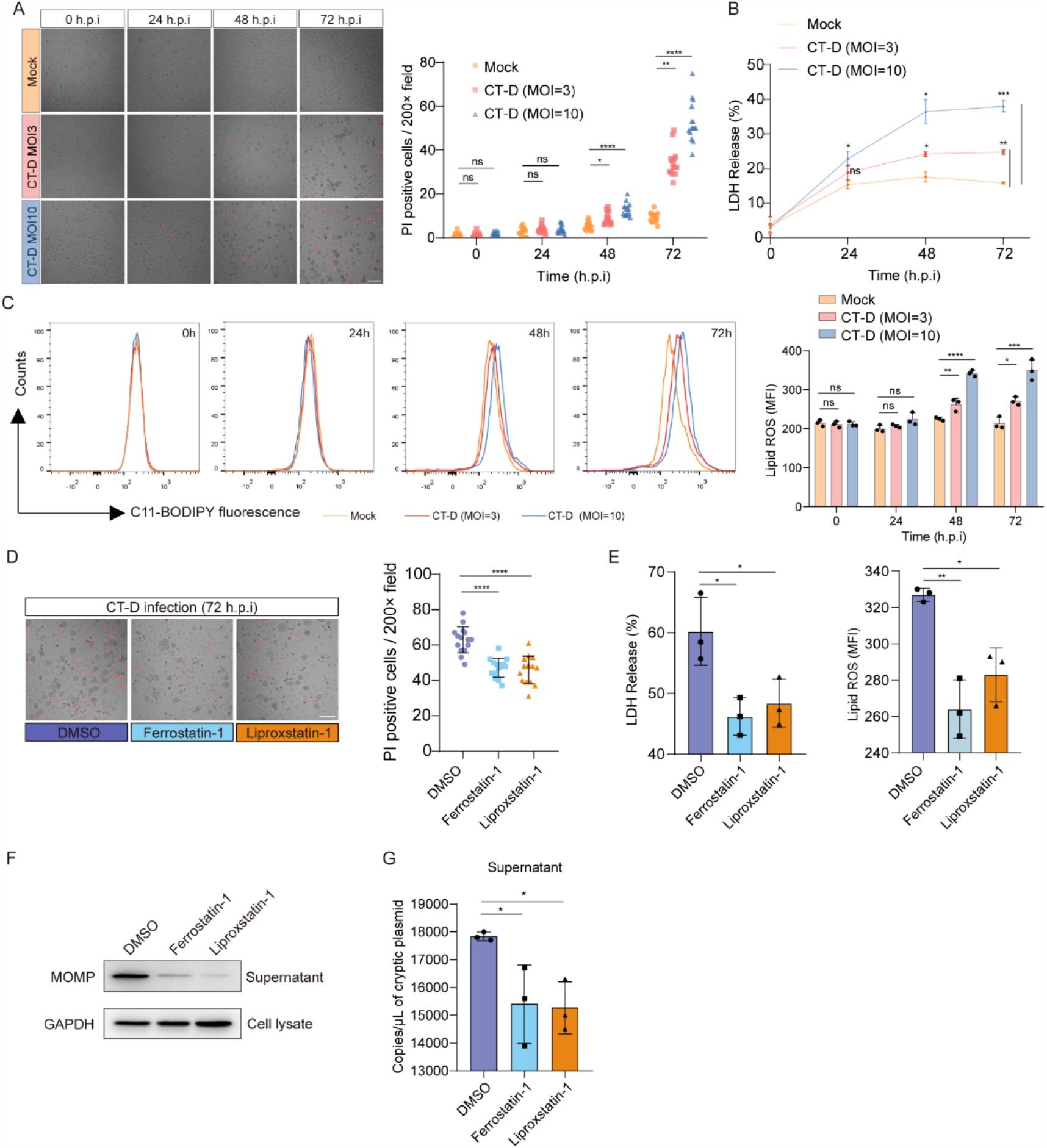
*Chlamydia* induces ferroptosis in host cells to facilitate the release of progeny. a) Host cell death induced by *Chlamydia trachomatis* serovar D (CT-D) at various multiplicities of infection (MOIs) was evaluated over a time course of 0-72 hours post-infection using a propidium iodide (PI) staining assay. PI-positive cells per 200 × field was counted for subsequent statistical analysis. The results demonstrated that CT-D infection induced necrotic cell death in host cells, particularly at the late stage of infection. Scale bars, 100 μm; b) At the late stage of infection, the release of lactate dehydrogenase (LDH) was significantly increased in CT-D infected cells compared to mock-infected cells, and this effect was observed across different MOIs; c) The level of lipid peroxidation in CT-D infected cells with different MOIs across the infection was assessed using C11-BODIPY staining throughout the course of infection. Accumulated lipid ROS was observed in CT-D infected cells at 48 h and 72 h post-infection; d) The cell death was measured by PI staining assay. PI-positive cells per 200X field was counted. Two ferroptosis inhibitors, ferrostatin-1(10 μM) and liproxstatin-1 (1μM) exhibited notable inhibition to ferroptosis driven by CT-D; e) LDH release and lipid ROS levels in CT-D infected cells were significantly reduced by ferrostatin-1(10 μM) and liproxstatin-1 (1μM); f) Ferrostatin-1 (10 μM) and liproxstatin-1 (1μM) blocked CT-D progeny release, as indicated by the presence of chlamydial MOMP in the supernatant; g) CT-D progeny release was restricted by ferrostatin-1 (10 μM) and liproxstatin-1 (1μM), as determined by quantitation of CT cryptic plasmid in the supernatant. Data are mean ± SD (n=3, except for A and D). *, < 0.05; **, < 0.01; ***, <0.001; ****, <0.0001; ns, not significant.

We also identified increased lactate dehydrogenase (LDH) release and production of lipid reactive oxygen species (ROS) in host cells infected with CT serovar L1 (CT-L1) and *Chlamydia muridarum* (CM), as illustrated in Supplementary Fig. S1a and S1b. These effects were observed in both HeLa-229 cells and mouse fibroblast cells (McCoy), indicating that this mechanism is independent of the cell type. Pharmacological inhibition of ferroptosis effectively reduced the cell death induced by CT-L1 and CM, as shown in Supplementary Fig. S1c and S1d. The capacity of inducing host cell ferroptosis across different *Chlamydia* strains or serovars underscores the conservation of this mechanism.

We proceeded to investigate whether the inhibition of ferroptosis could effectively attenuate the release of CT progeny. Immunoblotting of the major outer membrane protein (MOMP) or chlamydial HSP60 of CT or CM in the culture supernatant demonstrated that treatment with ferroptosis inhibitors led to a significant decrease in the release of progeny (Fig. 1f and Supplementary Fig. S1e). This observation was further confirmed by quantifying the copy number of chlamydial cryptic plasmids in the culture supernatant (Fig. 1g and Supplementary Fig. S1f). Collectively, these findings provide compelling evidence that *Chlamydia* strategically utilizes the host ferroptosis machinery to enhance its dissemination.

### Ferroptosis is induced by depletion of SLC7A11 and reduced levels of glutathione

To elucidate the mechanism by which *Chlamydia* manipulates ferroptosis, we assessed the involvement of two key cellular antioxidant systems, namely the System Xc−/glutathione (GSH)/GPx4 (23, 24) and FSP1/CoQ (25, 26) pathways, which play a critical role in defending against ferroptosis, during the late stage of CT infection. We observed a significant reduction in the abundances of SLC7A1 and GPx4 in cells infected with CT-D compared to mock-infected cells (Fig. 2a). However, we did not observe a significant change in the levels of FSP1 under these conditions. The substantial decrease in the abundance of SLC7A11 and GPx4 were confirmed in CT-L1 and CM infected host cells (Fig. 2b). SLC7A11 serves as a membrane antiporter, facilitating the uptake of cystine and enabling the subsequent synthesis of GSH, which collaborates with GPx4 in exerting anti-ferroptotic effects (Fig. 2c)(24). Therefore, we proceeded to investigate whether the level of intracellular GSH in infected cells was affected by the CT-induced deficiency of SLC7A11. As shown in Fig. 2d, CT-infected cells displayed a substantial decrease in GSH levels compared to mock-infected cells. These findings support the involvement of the SLC7A11-GSH-GPx4 axis in

**Fig. 2.**
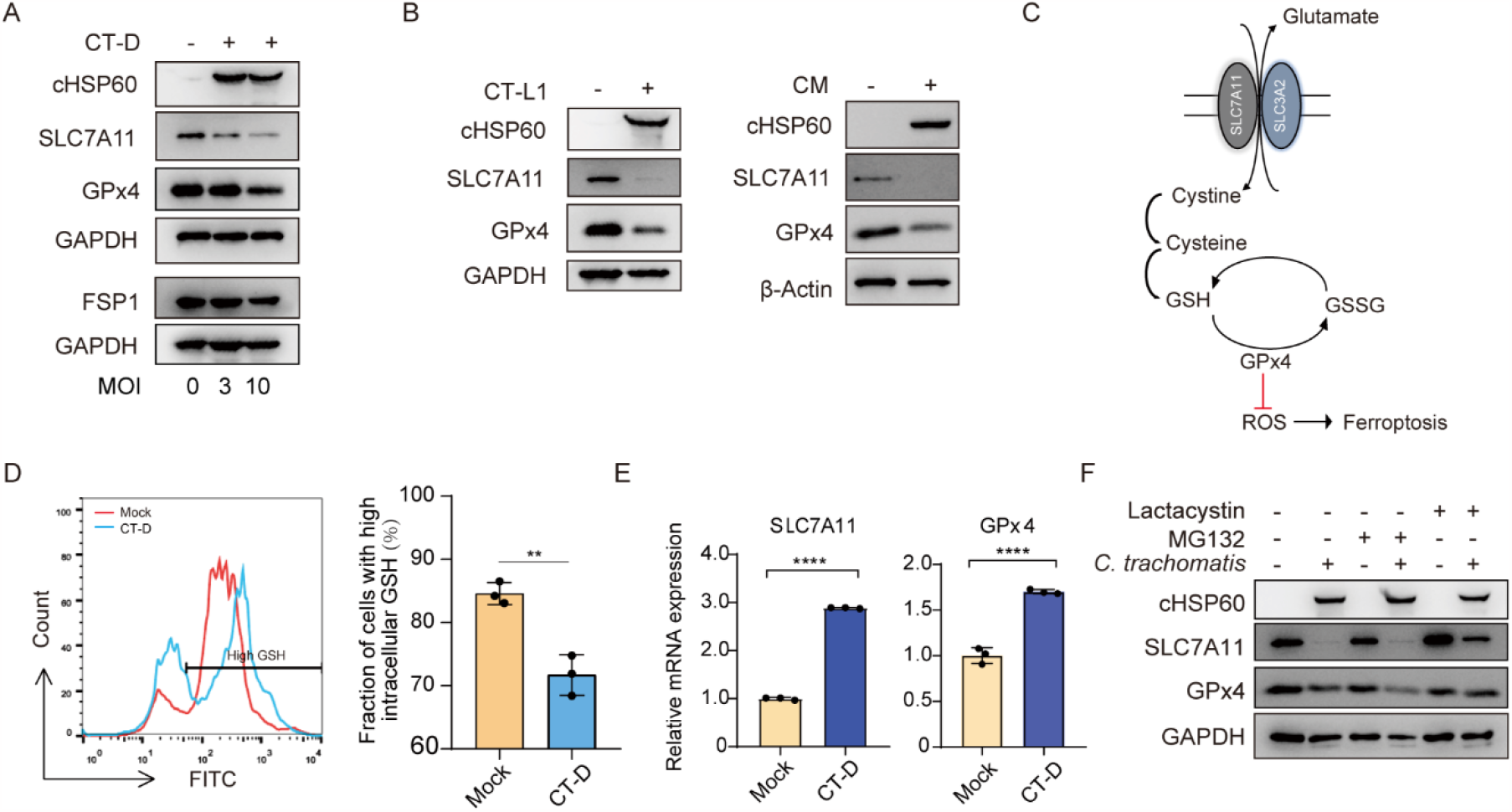
Ferroptosis is induced by depletion of SLC7A11 and reduced levels of glutathione. a) At 72 hours post-infection, cells infected with *Chlamydia trachomatis* serovar D (CT-D) showed a reduction in the protein levels of SLC7A11 and GPx4 compared to mock-infected cells; b) The deficiency of SLC7A11 and GPx4 was confirmed in *Chlamydia trachomatis* serovar L1 (CT-L1) and *Chlamydia muridarum* (CM)-infected cells at 72 h post-infection, compared to mock-infected cells; c) A schematic representation of the SLC7A11-GSH-GPx4 pathway in the modulation of ferroptosis; d) At 72 hours post-infection, a significant decrease in intracellular glutathione (GSH) levels was observed in CT-infected cells compared to mock-infected cells. This decrease in GSH levels suggests that CT infection leads to a deficiency in GSH levels; e) The expression of SLC7A11 and GPx4 was found to be up-regulated in CT-infected cells compared to mock-infected cells at 72 h post-infection; f) After a 6-hour pre-treatment with lactacystin (10 μM) or MG132 (10 μM), the decreased expression of SLC7A11 in CT-infected cells was restored by lactacystin treatment at 72 hours post-infection, whereas MG132 treatment did not exhibit the same rescue effect. Data are mean ± SD (n=3 for D and E). *, < 0.05; **, < 0.01; ***, <0.001; ****, <0.0001; ns, not significant.

#### *Chlamydia*-induced ferroptosis

We observed the mRNA expressions of SLC7A11 and GPx4 were upregulated in infected cells (Fig. 2e), which is consistent with the pattern of gene expression changes associated with ferroptosis (27). This suggests that *Chlamydia* did not alter the transcriptional regulation of SLC7A1 and GPx4 during this process. Hybiske *et al* (4) previously described that the lysis activity of CT-infected cells was relative to proteases activity. To assess the potential involvement of protease activity in the reduced abundances of SLC7A11 and GPx4, we treated the infected cells with two proteasomal inhibitors, MG132 and lactacystin. Interestingly, only treatment with lactacystin resulted in the restoration of SLC7A11 and GPx4 levels (Fig. 2f). Previous studies have demonstrated the specific inhibitory effect of lactacystin, but not MG132, on the protease activity of chlamydial protease-like activity factor (CPAF) (15, 28). Based on these findings, we proposed that CPAF may potentially play a role in mediating this process.

### CPAF-deficient *Chlamydia* fails to trigger ferroptosis and promote progeny release

CPAF, a potent virulence factor of *Chlamydia*, exerts multiple functions such as defending against host immunity, maintaining pathogen vacuole integrity, inhibiting apoptosis, and altering cell cycle, through its ability to cleave various host and chlamydial substrates (15-17, 29-31). To investigate whether CPAF participated in driving ferroptosis in the late stages of CT infection, we employed two transgenic organisms L2-17/mCherry (CPAF-deficient) and L2-17/CPAF(CPAF-complement) for determining the *Chlamydia*-driven ferroptotic events. Both transgenic organisms were obtained by transforming pGFP::SW2 plasmid expressing a wild-type allele of CPAF or mCherry into the CPAF-deficient strain L2-17 as described previously (19, 32).

Immunoblotting analysis confirmed the expression of CPAF in each transgenic strain, and notable differences in cell death morphology were observed between the infected cells of the two transgenic strains (Fig. 3a). A substantial decrease in LDH release was observed in cells infected with the CPAF-deficient strain L2-17/mCherry compared to the CPAF-complemented strain L2-17/CPAF (Fig. 3b). The presence of the GFP coding element on the pGFP::SW2 plasmid in both transgenic organisms hindered the practical application of BODIPY™ 581/591 C11 for lipid peroxide detection. As an alternative, we employed TfR1 as a specific ferroptosis marker to investigate the potential role of CPAF in triggering ferroptosis (33). The expression of TfR1 was significantly upregulated in cells infected with CPAF-expressing strains (L2-17/CPAF), whereas infection with the CPAF-deficient strain L2-17/mCherry showed similar expression levels to mock infection (Fig. 3c). In line with this, the decreased levels of SLC7A11 and GPx4 were correlated with the presence of CPAF. The CPAF-deficient strain CT-17/mCherry did not exhibit alterations in the protein levels of these anti-ferroptosis factors (Fig. 3c). Furthermore, the CPAF-deficient strain (CT-17/mCherry) exhibited impaired progeny release, as evidenced by reduced levels of chlamydial HSP60 and cryptic plasmid in the supernatant (Fig. 3d). Collectively, these findings strongly suggest that *Chlamydia*-induced ferroptosis and subsequent dissemination of progeny are dependent on the presence of CPAF.

**Fig. 3.**
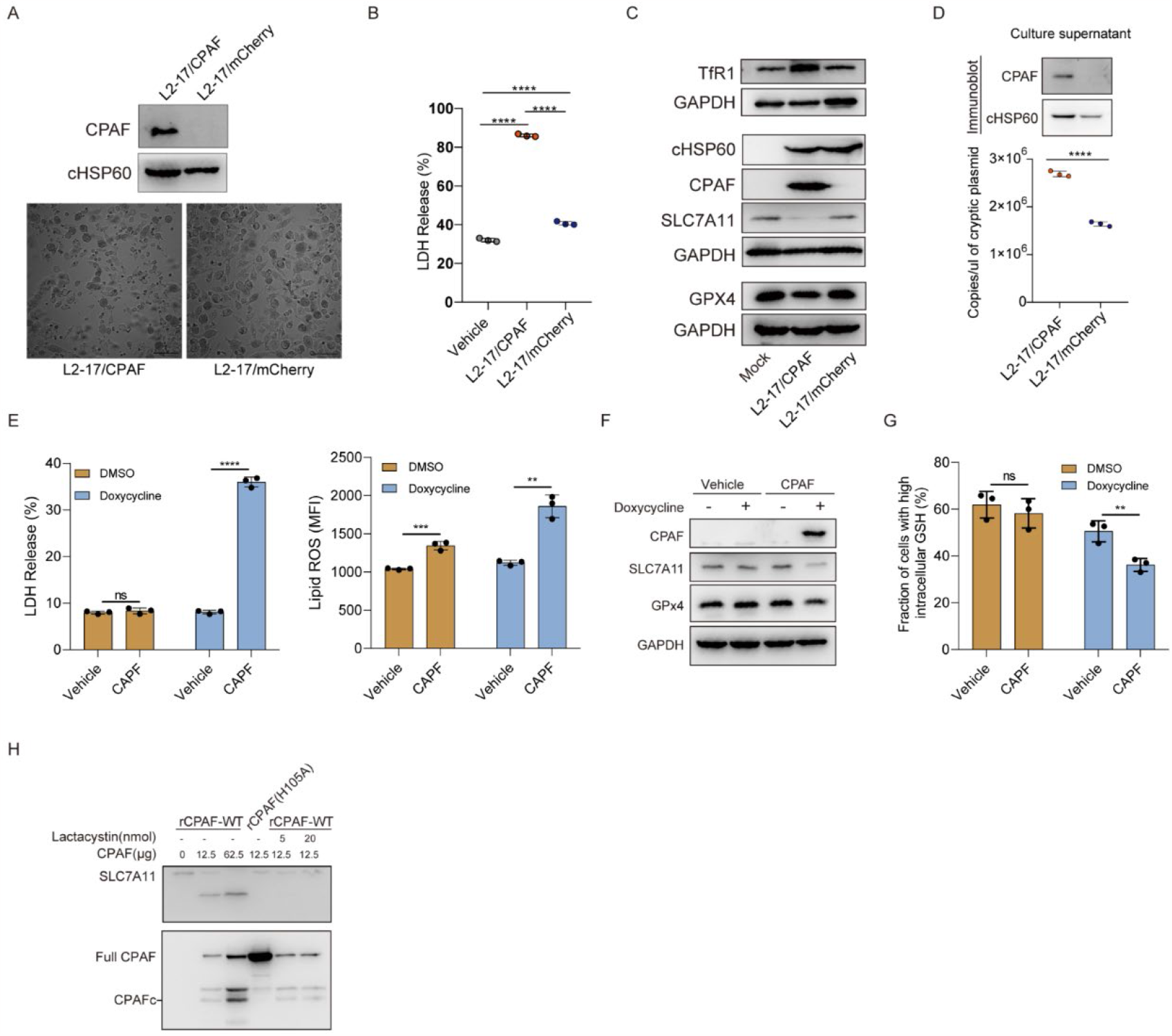
CPAF directly degrades host cell protein SLC7A11 and facilitates *Chlamydia trachomatis* dissemination. a) At 72 hours post-infection, the CPAF-deficient strain (L2-17/mCherry) exhibited fewer morphological changes indicative of cell death compared to the CPAF-sufficient strain (L2-17/CPAF). The absence of CPAF in the L2-17/mCherry strain was confirmed through western blotting; b) The release of lactate dehydrogenase (LDH) from cells infected with the CPAF-deficient strain (L2-17/mCherry) was significantly decreased compared to cells infected with the CPAF-sufficient strain (L2-17/CPAF) at 72 hours post-infection; c) At 72 hours post-infection, cells infected with the L2-17/mCherry strain did not show the upregulation of TfR1 and the decrease in the abundance of SLC7A11 compared to mock-infected cells; d) CT lacking CPAF showed impaired release of progeny at 72 hours post-infection, as indicated by decreased levels of chlamydial HSP60 and cryptic plasmid in the supernatant, as confirmed by both western blotting and quantitative qPCR analyses; e) The induction of CPAF expression in HeLa-229 cells resulted in elevated levels of LDH release and increased accumulation of lipid ROS at 72 h post-infection; f) The expression of CPAF and the degradation of SLC7A11 and GPx4 were confirmed in HeLa-229 cells after a 24-hour treatment with doxycycline, as determined by western blotting; g) After 24 hours of treatment with doxycycline, the intracellular glutathione (GSH) levels were found to be decreased in HeLa-229 cells; h) After a 30-minute incubation, recombinant wild-type CPAF (rCPAF), but not the H105A mutant, directly degraded SLC7A11 in cell lysates. Notably, an activated CPAFc fragment was observed in the wild-type CPAF group, while it was absent in the CPAF (H105A) group. The prototypic degradation of recombinant wild-type CPAF could be blocked by lactacystin. Data are mean ± SD (n=3 for B, D, E and G). *, < 0.05; **, < 0.01; ***, <0.001; ****, <0.0001; ns, not significant.

To investigate whether CPAF alone could induce ferroptotic events, we developed a eukaryotic HeLa-229 cell line with a tetracycline (Tet) inducible expression system for CPAF. Following a 24-hour treatment with doxycycline, we observed increased LDH release and lipid ROS levels in the CPAF-expressed cells (Fig. 3e), along with decreased abundances of SLC7A11 and GPx4 (Fig. 3f). Additionally, the cellular GSH level, downstream of SLC7A11, was significantly inhibited in the CPAF-expressed cells compared to controls (Fig. 3g). These findings suggest that CPAF alone is sufficient to induce ferroptotic events.

### CPAF targets SLC7A11 but not GPx4 for proteolytic degradation

To investigate whether CPAF can directly cleave SLC7A11 and GPx4, we expressed recombinant CPAF (rCPAF) and a mutant form of CPAF (H105A) that resulted in a defective proteolytic activity (28, 32). To inhibit the proteolytic activity of rCPAF, lactacystin was employed. As shown in Fig. 3h, rCPAF underwent autocatalytic cleavage, resulting in the generation of activated CPAFc fragments. In contrast, the mutant rCPAF(H105A) did not produce the activated CPAFc fragment. SLC7A11 was degraded by incubating with rCPAF but not rCPAF(H105A) or rCPAF combined with lactacystin treatment (Fig. 3h). Notably, rCPAF didn’t degrade GPx4 in vitro (Supplementary Fig. S2).

### Pharmacological inhibition of ferroptosis restricted *Chlamydia* infection

Pharmacological therapeutic strategies targeting ferroptosis hold promise for cancer immunotherapy and cardiomyopathy protection (34, 35). Vitamin E is known to function as a lipid-soluble antioxidant that prevents the propagation of lipid peroxidation (36, 37). Considering the beneficial role of ferroptosis in CT dissemination, we investigated the therapeutic potential of vitamin E in suppressing CT infection, both in vitro and in vivo. Due to the liposoluble nature of vitamin E, we utilized trolox, an analogue of vitamin E, to evaluate its effectiveness in vitro. The results showed that trolox successfully reduced the levels of LDH release and lipid ROS accumulation induced by CM in McCoy Cells (Fig. 4a). Additionally, trolox treatment inhibited the release of CM progeny, as evidenced by decreased levels of chlamydial HSP60 and reduced copy numbers of CM Nigg II plasmid in the supernatant (Fig. 4b).

**Fig. 4.**
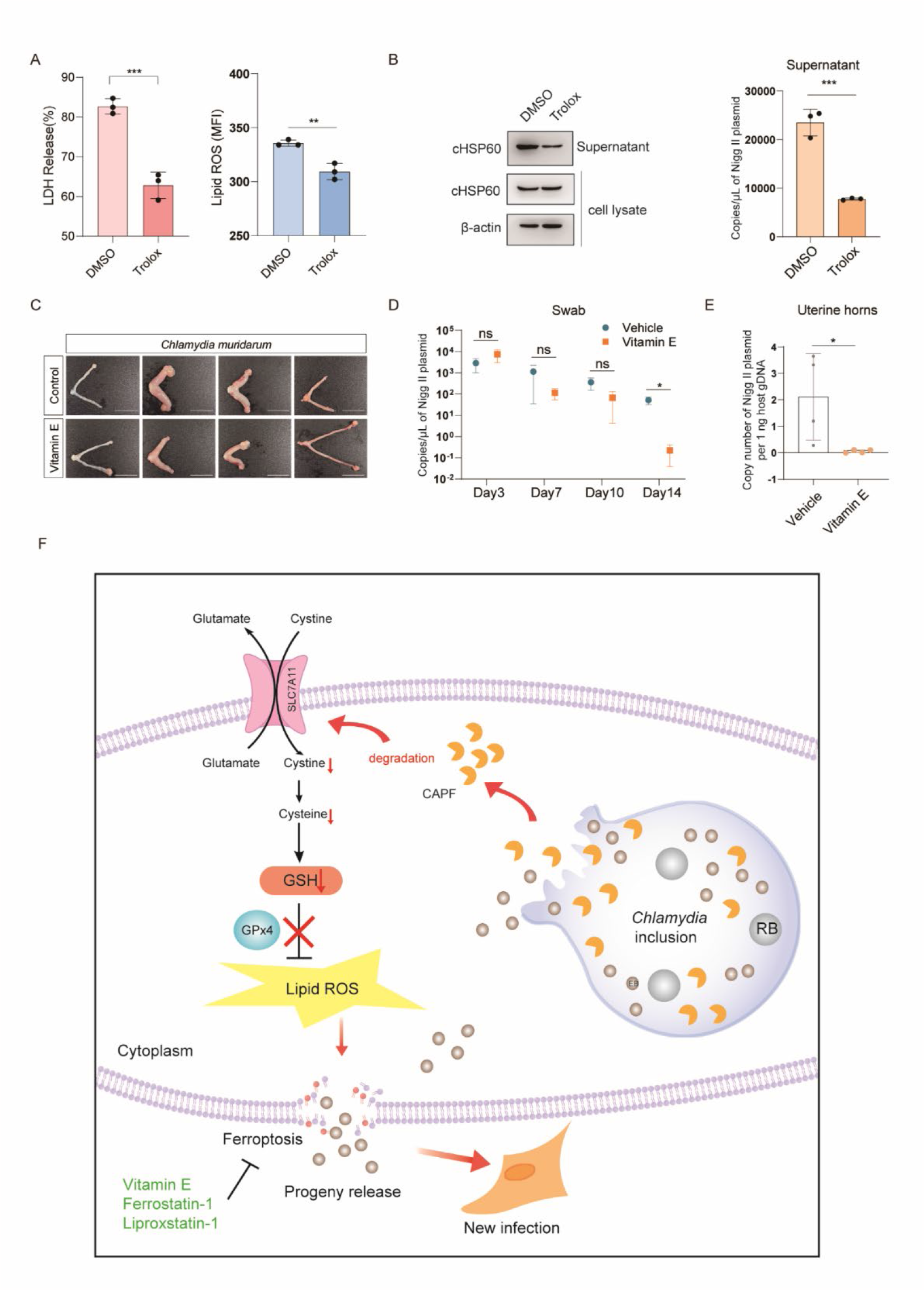
Pharmacological inhibition of ferroptosis restricted *Chlamydia* infection. a) Treatment with trolox, an analogue of vitamin E, resulted in a decrease in lactate dehydrogenase (LDH) release and accumulation of lipid reactive oxygen species (ROS) in McCoy cells infected with *Chlamydia muridarum* (CM); b) The release of CM progeny was inhibited by the administration of Trolox; c) Mice infected with CM and treated with vitamin E exhibited reduced dilation of the uterine horns compared to untreated mice (n=4). Scale bars, 1 cm; d) The copy numbers of CM plasmid in vaginal swabs collected from mice treated with vitamin E were significantly lower 14 days post-infection compared to the vehicle-treated group; e) Vitamin E treatment led to a remarkable reduction in *Chlamydia* burdens in the uterine horns of infected mice compared to the vehicle-treated group; f) A schematic diagram depicts the key findings of this study, highlighting the direct degradation of the host protein SLC7A11 by CPAF, leading to the induction of ferroptosis and facilitating the dissemination of *Chlamydia*. Data are mean ± SD (n=3 for A and B; n=4 for D and E). *, < 0.05; **, < 0.01; ***, <0.001; ****, <0.0001; ns, not significant.

We established a CM-infected mouse model and evaluated *Chlamydia* loads in vaginal swabs at various time points (day 3, 7, 10, and 14) post-infection, with and without vitamin E treatment. After a 14-day infection period, the mice were euthanized, and their uterine horns were examined. CM-infected mice treated with vitamin E exhibited reduced dilation of the uterine horns (Fig. 4c). Furthermore, the copy numbers of chlamydial Nigg II plasmid in vaginal swabs collected from vitamin E-treated mice were significantly lower compared to the vehicle groups, indicating the inhibitory effect of vitamin E on *Chlamydia* infection (Fig. 4d). Additionally, when we analyzed the *Chlamydia* loads in the uterine horns of infected mice, we observed that vitamin E treatment significantly reduced the *Chlamydia* burdens compared to the vehicle treatment (Fig. 4e). These findings suggest that vitamin E treatment effectively restricts *Chlamydia* dissemination to the upper-genital tract, highlighting its potential as a therapeutic intervention for *Chlamydia* infection.

## Discussion

*Chlamydia*, with its successful strategies for dissemination and propagation within the host, contributes significantly to reproductive health complications (1, 2). The lysis mechanism is an essential dissemination strategy of CT within the host (4). However, the underlying molecular mechanisms driving this process remain unexplored. While programmed cell death is recognized as a defensive strategy against pathogen infection, pathogens often capitalize on host mechanisms to their advantage during the ongoing host-pathogen interaction (5). In this study, we unveiled a previously unexplored association between ferroptosis and CT, elucidating its involvement in the lysis mechanism and facilitation of *Chlamydia* progeny release (Fig. 4f).

Emerging evidence suggests that ferroptosis, despite being relatively understudied in the context of infectious diseases, plays a significant role in promoting pathogen replication, dissemination, and pathogenesis (7-12). Pathogens, especially viruses, manipulate the lipid ROS process by targeting the GPX4-dependent antioxidant pathway through transcriptional regulation, disabling the SLC7A11-GSH-GPX4 axis and promoting pathogen propagation (7, 12). For example, *Hepatitis B virus* employs HBx to induce the enrichment of EZH2 and H3K27me3 in the SLC7A11 promoter, leading to the suppression of SLC7A11 gene expression (12).

*Mycobacterium tuberculosis* employs the secreted effector protein PtpA to interact with RanGTP, enabling its entry into the host cell nucleus, where it facilitates PRMT6-mediated H3R2me2a on the GPX4 promoter, resulting in the suppression of GPx4 mRNA expression (11). In contrast to previous reports, our study revealed that *Chlamydia* induces host cell ferroptosis through a direct proteolytic degradation of SLC7A11 instead of the commonly reported epigenetic transcription regulation. The deficiency of SLC7A11 in host cells at the late-stage infection was observed when challenged with various CT serovars as well as *Chlamydia muridarum*. Based on existing evidence (7, 11, 12) and our finding, the classical antioxidant system (SLC7A11-GSH-GPx4 pathway) appears to be more frequently targeted by intracellular pathogens compared to other antioxidant systems, indicating a conserved evolutionary pathway of intracellular pathogens in their long-term battle with the host.

CPAF, a serine chlamydial protease, harbors multiple functions in promoting CT survival by cleaving a broad of host and chlamydial substrates (15-17, 29-31). Despite previous proposals of CPAF’s involvement in CT dissemination (18, 20, 21), the specific molecular mechanism underlying its role is still not well understood. In the current study, we identified CPAF induces host cell ferroptosis through the direct proteolytic degradation of SLC7A11. Although previous studies have reported some targets were identified as substrates of CPAF as a result of post-lysis protease effects, cell lysis with 8M urea can help resolve this misdirection (38). The degradation of SLC7A11 was confirmed by western blot analysis with cell lysis conducted via urea lysis (Supplementary Fig.S3). Of significant importance is the observation that depletion of SLC7A11 led to a discernible reduction in intracellular GSH levels within CT-infected cells. This observation serves to suggest that the degradation of SLC7A11 occurred not as a consequence of post-lysis protease effects mediated by CPAF but rather during the course of infection. Moreover, the expression of CPAF in eukaryotic cells alone is sufficient to induce necrotic cell death, which supports the findings reported by Paschen *et al* (39). The ferroptosis phenotype and associated molecular changes observed in mammalian cells expressing CPAF are in line with observations in CT strain, providing further confirmation of CPAF’s role in triggering ferroptosis in host cells. Our results also demonstrate that GPx4 was suppressed in CT-infected or CPAF-expressed cells, but it was not degraded by recombinant CPAF in vitro, indicating that GPx4 is not a substrate of CPAF. In light of this finding, a possible explanation for the reduced abundance of GPx4 in infected or CPAF-expressed cells is that the degradation of SLC7A11 by CPAF limits selenium uptake and selenocysteine synthesis, ultimately leading to the inhibition of selenoprotein GPx4 synthesis (40).

Yang et al. found that the deficiency of CPAF impaired chlamydial survival in the mouse lower genital tract (19). On one hand, CPAF acts as an effector that paralyzes polymorphonuclear neutrophils upon *Chlamydia* exposure to host immunosurveillance, evading the host’s innate immune response by suppressing the oxidative burst, interfering with neutrophil activation, and cleaving FPR2(17). On the other hand, the absence of CPAF in *Chlamydia* fails to induce ferroptosis in host cells, resulting in impaired release of *Chlamydia* progeny, ultimately leading to compromised dissemination and reduced survival.

Pharmacological modulation of ferroptosis has been established that is beneficial for cancer suppression (34) and preventing ischemic organ injuries (35), suggesting the potential value of modulation of this cell death modality as a therapeutic avenue. Our study indicates that inhibition of ferroptosis could depress the *Chlamydia* progeny release in vitro and restrict the *Chlamydia* infection in the mouse infection model. This finding offers an explanation as to why the administration of vitamin E facilitates the recovery of lambs afflicted with intratracheal *Chlamydia* infection (14). Given the ferroptosis-specific activities of CT, host-directed therapies targeting ferroptosis used in conjunction with suitable antibiotics are a promising strategy for the treatment of CT infection.

## Materials and Methods

### Cell culture and *Chlamydia* infection

HeLa-229 cells (CCTCC Cat# GDC0335) and McCoy cells (ATCC Cat# CRL-1696) were cultured in high-glucose Dulbecco’s Modified Eagle’s Medium (DMEM) (Thermo Fisher Scientific, Grand Island, NY, USA; 11995065) supplemented with 10% fetal bovine serum (Gibco; 10099141C) at 37°C and 5% CO_2_. Prior to experimentation, the exclusion of Mycoplasma contamination was confirmed using a commercial kit (Biological Industries, Beit-Haemek, Israel; 20-700-20).

To establish the infected cell model, cells were challenged to *Chlamydia trachomatis* serovars D, L1, L2, and *Chlamydia muridarum* at specific multiplicities of infection (MOIs). In the lipid peroxidation inhibition experiment, ferrostatin-1 (10 μM; MedChemExpress; Shanghai, China; HY-100579), liproxstatin-1 (1 μM; Cayman Chemical, Ann Arbor, MI, USA; 17730-5), and trolox (3.2 mM; MedChemExpress; Shanghai, China; HY-101445) were added after the infection process. To assess proteolytic degradation during the late stage of infection, lactacystin (10 μM; Adipogen Life Sciences; San Diego, CA, USA; AG-CN2-0442-C100) and MG132 (10 μM; MedChemExpress; Shanghai, China; HY-13259) were added to infected cells 6 hours before determining the protein levels at 72 hours post-infection (h.p.i). The cell culture was maintained with 1 μg/ml of cycloheximide (MedChemExpress; Shanghai, China; HY-12320) throughout the experiment. Three independent experiments were performed for analysis.

### Cell death assay

Necrotic cell death was evaluated by staining cells with 500 nM of propidium iodide (PI; Thermo Fisher Scientific, Waltham, MA, USA; P3566). Following a 5-minute staining period, PI-positive cells were visualized using confocal microscopy (Nikon, Tokyo, Japan; Model A1R; RRID:SCR_020317). Cell viability was evaluated by measuring lactate dehydrogenase (LDH) release. The cell culture supernatant was collected and subjected to analysis using the CytoTox 96 assay (Promega, Madison, WI, USA; G1780) according to the manufacturer’s instructions. Positive control wells were included to determine the maximum LDH release and calculate the percentage of LDH release. Three independent experiments were performed for analysis.

### Analysis of lipid peroxide production

Lipid peroxide production was assessed using the lipophilic fluorescent dye C11-BODIPY 581/591 (Thermo Fisher Scientific, Waltham, MA, USA; D3861). Following treatment with the test compounds, the cell culture supernatant was replaced with 1 ml of fresh DMEM containing 5 μL of BODIPY, and incubated for 30 minutes at 37°C. The cells were then harvested using TrypLE Express Enzyme (Thermo Fisher Scientific; Waltham, MA, USA; 12604021) after three washes with phosphate-buffered saline (PBS). Subsequently, the cells were resuspended in fresh PBS for flow cytometry analysis using a BD FACSCelesta cell analyzer (BD Biosciences, Franklin Lakes, NJ, USA). The oxidation of the polyunsaturated butadienyl portion of the dye induces a shift in the fluorescence emission peak from 590 nm to 510 nm. Three independent experiments were performed for analysis.

### Western blotting

For western blot analysis, cultured cells or the culture supernatant was lysed using radioimmunoprecipitation assay (RIPA) buffer (Thermo Fisher Scientific, Waltham, MA, USA; 89900) supplemented with a protease and phosphatase inhibitor cocktail (Invitrogen; Thermo Fisher Scientific, Waltham, MA, USA; 78440) or 8M urea buffer (38). Subsequently, the samples were incubated with reducing sodium dodecyl sulfate-polyacrylamide electrophoresis loading buffer (CWBio, Beijing, China; CW0027) at 100? for 10 minutes. Antibodies against specific proteins were used as follows: GPx4 (1:1000, Abcam Cat# ab125066), SLC7A11 (1:1000, Abcam Cat# ab175186), major outer membrane protein (MOMP; 1:1000, Abcam Cat# ab20881), and glyceraldehyde 3-phosphate dehydrogenase (GAPDH; 1:5000, Abcam Cat# ab181602); fibroblast-specific protein 1 (FSP1; 1:500, Santa Cruz Biotechnology Cat# sc-377120), TfR1(1:500, Santa Cruz Biotechnology Cat# sc-32272), Vimentin (1:1000, Santa Cruz Biotechnology Cat# sc-6260); ERK1/2 (1:1000, Bio-Rad Cat# MCA4695T), horseradish peroxidase-conjugated anti-rabbit (1:5000, Cell Signaling Technology Cat# 7074S), and antimouse (1:5000, Cell Signaling Technology Cat# 7076S) IgG; mouse monoclonal antibody clones 100a against a 35 kDa activated CPAFc (kindly provided by Dr. Lingli Tang, 1:50 dilution). Blots were imaged using a ChemiDoc imaging system (v2.4.0.03; Bio-Rad; Hercules, CA, USA). Three independent experiments were performed for analysis.

### Assessment of intracellular GSH

Intracellular glutathione (GSH) levels were measured using the Glutathione (GSH) kit (Abcam; Cambridge, MA, USA; ab112132). HeLa-229 cells were collected and washed with PBS, followed by incubation with a green dye at 37°C for 30 minutes. The fluorescence intensity of the green dye was detected using a flow cytometer at the FITC channel. Three independent experiments were performed for analysis.

### Quantitative real-time PCR of *Chlamydia* from culture supernatant or cell lysates

CT DNA was extracted from the culture supernatant or cell lysates using the TIANamp Genomic DNA kit (Tiangen Biotech, Beijing, China; DP304). Quantitative PCR analysis targeting the CT cryptic plasmid was performed using TaqMan Gene Expression Master (Thermo Fisher Scientific, Waltham, MA, USA; 4369016) with a standard curve ranging from 1.0×10^2^ to 1.0×10^8^ copies/μL. The plasmids, primers, and probes used in the assay were synthesized by Sangon Biotech (Shanghai, China). The specific primers and probes used for quantifying CT were as follows: (forward) 5’-ATTTTGGCCGCTAGAAAAGGC-3’, (reverse) 5’-CGGAACACATGATGCGAAGT-3’, and probe 5’-ROX-CGAACTCATCGGCGATAAGG-3’-BHQ2 targeting the cryptic plasmid. Three independent experiments were performed for analysis.

### Relative mRNA expression of SLC7A11 and GPx4

Total RNA was extracted from *Chlamydia*-infected HeLa-229 cells at 72 hours post-infection using the TRIzol reagent (Thermo Fisher Scientific, Waltham, MA, USA; 15596018). First-strand cDNA was prepared using the Prime Script RT Reagent Kit (Takara Bio, Shiga, Japan; RR047B) according to the manufacturer’s instructions. The relative mRNA expression was measured by TB Green Premix Ex Taq II (Takara Bio, Shiga, Japan; RR82WR) according to the manufacturer’s instructions. The primers used for SLC7A11, GPx4, and GAPDH were synthesized by Sangon Biotech (Shanghai, China) and had the following sequences: SLC7A11 (forward) 5’-CCTCTGCCAGCTGTTATTGTT-3’, (reverse) 5’-CCTGGCAAAACTGAGGAAAT-3’; GPx4 (forward) 5’-GCAACCAGTTTGGGAGGCAGGAG-3’, (reverse) 5’-CCTCCATGGGACCATAGCGCTTC-3’; GAPDH (forward) 5’-ATGGCACCGTCAAGGCTGAG-3’, (reverse) 5’-GCAGTGATGGCATGGACTGT -3’. Three independent experiments were performed for analysis.

### Inducible CPAF expression in HeLa-229 cells

The coding sequence of *Chlamydia trachomatis* CPAF (amino acid residues 18-601) was cloned into the pLVX-TetOne-Puro plasmid (P1686) obtained from MIAOLING Biology, Wuhan, China. The pLVX-TetOne-CPAF-puro plasmid, along with packaging plasmids, was co-transfected into HEK293T cells (CCTCC; Cat# GDC0187) to generate infectious lentivirus particles. The lentivirus particles were harvested from the cell culture supernatant. HeLa-229 cells were then transduced with the lentivirus in the presence of polybrene to facilitate efficient viral entry. To establish inducible CPAF expression in HeLa-229 cells, puromycin was added to the cell culture media to selectively pressure and generate stable cell lines. Similarly, transduction with the empty pLVX-TetOne-puro vector (vehicle) was performed as a control. The vehicle and CPAF-inducible expressed HeLa-229 cells were cultured overnight. CPAF expression was induced by treating the cells with 1 μg/mL doxycycline for 24 hours in preparation for subsequent experiments.

### Cell-free Degradation Assay

To measure CPAF degradation of SLC7A11, a cell-free degradation assay was performed. Fusion proteins were generated for the assay by cloning chlamydial DNA sequences encoding CPAF into a pET30a vector (constructed by Genscript Biotech), resulting in fusion proteins with a hexahistidine (His6) tag as the partner. HeLa-229 cells were lysed on ice for 10 minutes using RIPA buffer. The cleared lysates, which served as the source of host protein substrates, were obtained by centrifugation at 14,000×g for 15 minutes at 4°C. Recombinant wild-type CPAF or mutant CPAF (H105A) at various concentrations were pre-incubated with the HeLa-229 cell lysates at room temperature for 30 minutes, in the presence or absence of the proteasome inhibitor lactacystin (Adipogen Life Sciences; San Diego, CA, USA; AG-CN2-0442-C100) at the indicated concentration.

### Murine model of *Chlamydia muridarum* infection

All animal experiments conducted in this study were approved by the Animal Welfare Committee of South China Agricultural University (2022G007) and were conducted in accordance with the regulations and guidelines of the institution. Female mice aged four weeks received subcutaneous administration of 2.5 mg of progesterone (Baiyunshan Pharmaceutical General Factory, Guangzhou, China; H44020229) at 10 and 3 days prior to *Chlamydia muridarum* infection. On the day of infection, mice were intravaginally infected twice with 9×10^5^ inclusion forming units (IFU) in 10μL sucrose-phosphate-glutamate buffer (SPG, pH 7.5) buffer. After the second infection, the control group was administered oil, while the experimental group received oral administration of vitamin E (Sigma-Aldrich; Burlington, MA, USA; 258024) (500 mg/kg of body weight, dissolved in corn oil). The control and experimental groups were given the respective treatments again at 7 days post-infection (d.p.i). To monitor the infectious burden, vaginal swabs were collected at 3, 7, 10, and 14 d.p.i. On the 14th day of infection, the mice were sacrificed, and the uterine horn was collected for DNA extraction.

DNA extraction from swabs and tissue was performed using the DNeasy Blood & Tissue kit (QIAGEN; Hilden, Germany; 69506). Quantification of chlamydia DNA in the extracts was conducted by qPCR using primers and probes targeting the *Chlamydia muridarum* Nigg II plasmid. Quantitative PCR based on the Nigg II plasmid was performed against a standard curve ranging from 1.0×10^1^ to 1.0×10^7^ copies/μL, using TaqMan Gene Expression Master (Thermo Fisher Scientific, Waltham, MA, USA; 4369016). The specific primers and probe were as follows: (forward) 5’-GTCATTCTGTTTAAAAATCTAGTCAAA-3’, (reverse) 5’-TTCATGGACAGAAGGCACC-3’, and probe 5’-FAM-GATATAACTAGCTGCACGAACTTG-3’-BHQ1 for the Nigg II plasmid. Mouse β-actin was used as a reference gene to normalize the input of experimental uterine horns by comparing it to a known concentration of gDNA from uterine horns. The primers of mouse β-actin were as follows: (forward) 5’-GTGCTATGTTGCTCTAGACTTCG -3’, (reverse) 5’-ATGCCACAGGATTCCATACC -3’. All plasmids, primers, and probes were synthesized by Sangon Biotech (Shanghai, China).

### Statistical analysis

Data analysis and visualization were performed using Prism 8 software (GraphPad, La Jolla, CA, USA). Quantitative data were presented as mean ± standard deviation and were assessed for normality using the Shapiro-Wilk test. Multiple comparisons were analyzed using one-way ANOVA, while two-group comparisons were performed using Student’s *t*-test. Statistical significance was determined at a threshold of *P*<0.05.

## Acknowledgments

We thank Dr. Lingli Tang from Second Xiangya Hospital of Central South University for providing the L2-17/mCherry, L2-17/CPAF strains and CPAF antibody (100a). We express our gratitude to Dr. Xiaomian Lin and other colleagues at the Dermatology Hospital of Southern Medical University for their valuable assistance and support throughout this study.

## Author Contributions

W.C. conceptualized the study. W.C., X.S., Y.P., Y.X. designed the experiments. W.C., X.S., Y.P., Y.X., L.Z., X.Y., Q.X., X.Y., H.Z. performed most of the experiments, data analysis, and data interpretation. Z.F., B.Z., W.Z., H.Z. are the study supervisor. W.C., X.S., Y.P., Y.X. drafted the manuscript. All authors contributed to editing, reviewing of the manuscript, and approved the final version of the manuscript.

## References

1. S. Menon et al., Human and Pathogen Factors Associated with Chlamydia trachomatis-Related Infertility in Women. Clinical microbiology reviews 28, 969–985 (2015).

2. C. D. J. den Heijer et al., Chlamydia trachomatis and the Risk of Pelvic Inflammatory Disease, Ectopic Pregnancy, and Female Infertility: A Retrospective Cohort Study Among Primary Care Patients. Clinical infectious diseases : an official publication of the Infectious Diseases Society of America 69, 1517–1525 (2019).

3. K. Stelzner, N. Vollmuth, T. Rudel, Intracellular lifestyle of Chlamydia trachomatis and host-pathogen interactions. Nature reviews. Microbiology 10.1038/s41579-023-00860-y (2023).

4. K. Hybiske, R. S. Stephens, Mechanisms of host cell exit by the intracellular bacterium Chlamydia. Proc Natl Acad Sci U S A 104, 11430–11435 (2007).

5. I. Jorgensen, M. Rayamajhi, E. A. Miao, Programmed cell death as a defence against infection. Nat Rev Immunol 17, 151–164 (2017).

6. S. J. Dixon et al., Ferroptosis: an iron-dependent form of nonapoptotic cell death. Cell 149, 1060–1072 (2012).

7. J. Gao et al., When ferroptosis meets pathogenic infections. Trends in microbiology 10.1016/j.tim.2022.11.006 (2022).

8. E. P. Amaral et al., A major role for ferroptosis in Mycobacterium tuberculosis-induced cell death and tissue necrosis. The Journal of experimental medicine 216, 556–570 (2019).

9. H. H. Dar et al., Pseudomonas aeruginosa utilizes host polyunsaturated phosphatidylethanolamines to trigger theft-ferroptosis in bronchial epithelium. The Journal of clinical investigation 128, 4639–4653 (2018).

10. D. Yamane et al., FADS2-dependent fatty acid desaturation dictates cellular sensitivity to ferroptosis and permissiveness for hepatitis C virus replication. Cell chemical biology 29, 799–810.e794 (2022).

11. L. Qiang et al., A mycobacterial effector promotes ferroptosis-dependent pathogenicity and dissemination. Nature communications 14, 1430 (2023).

12. G. Z. Liu, X. W. Xu, S. H. Tao, M. J. Gao, Z. H. Hou, HBx facilitates ferroptosis in acute liver failure via EZH2 mediated SLC7A11 suppression. Journal of biomedical science 28, 67 (2021).

13. A. A. Azenabor, J. B. Mahony, Generation of reactive oxygen species and formation and membrane lipid peroxides in cells infected with Chlamydia trachomatis. International journal of infectious diseases : IJID : official publication of the International Society for Infectious Diseases 4, 46–50 (2000).

14. L. C. Stephens, A. E. McChesney, C. F. Nockels, Improved recovery of vitamin E-treated lambs that have been experimentally infected with intratracheal Chlamydia. The British veterinary journal 135, 291–293 (1979).

15. I. Jorgensen et al., The Chlamydia protease CPAF regulates host and bacterial proteins to maintain pathogen vacuole integrity and promote virulence. Cell host & microbe 10, 21–32 (2011).

16. G. Zhong, P. Fan, H. Ji, F. Dong, Y. Huang, Identification of a chlamydial protease-like activity factor responsible for the degradation of host transcription factors. The Journal of experimental medicine 193, 935–942 (2001).

17. K. Rajeeve, S. Das, B. K. Prusty, T. Rudel, Chlamydia trachomatis paralyses neutrophils to evade the host innate immune response. Nature microbiology 3, 824–835 (2018).

18. E. A. Snavely et al., Reassessing the role of the secreted protease CPAF in Chlamydia trachomatis infection through genetic approaches. Pathogens and disease 71, 336–351 (2014).

19. Z. Yang et al., The Chlamydia-Secreted Protease CPAF Promotes Chlamydial Survival in the Mouse Lower Genital Tract. Infection and immunity 84, 2697–2702 (2016).

20. Y. Kumar, R. H. Valdivia, Actin and intermediate filaments stabilize the Chlamydia trachomatis vacuole by forming dynamic structural scaffolds. Cell host & microbe 4, 159–169 (2008).

21. C. Yang et al., Chlamydial Lytic Exit from Host Cells Is Plasmid Regulated. mBio 6, e01648–01615 (2015).

22. O. Zilka et al., On the Mechanism of Cytoprotection by Ferrostatin-1 and Liproxstatin-1 and the Role of Lipid Peroxidation in Ferroptotic Cell Death. ACS central science 3, 232–243 (2017).

23. W. S. Yang et al., Regulation of ferroptotic cancer cell death by GPX4. Cell 156, 317–331 (2014).

24. R. Yan et al., The structure of erastin-bound xCT-4F2hc complex reveals molecular mechanisms underlying erastin-induced ferroptosis. Cell research 32, 687–690 (2022).

25. K. Bersuker et al., The CoQ oxidoreductase FSP1 acts parallel to GPX4 to inhibit ferroptosis. Nature 575, 688–692 (2019).

26. S. Doll et al., FSP1 is a glutathione-independent ferroptosis suppressor. Nature 575, 693–698 (2019).

27. B. R. Stockwell, Ferroptosis turns 10: Emerging mechanisms, physiological functions, and therapeutic applications. Cell 185, 2401–2421 (2022).

28. Z. Huang et al., Structural basis for activation and inhibition of the secreted chlamydia protease CPAF. Cell host & microbe 4, 529–542 (2008).

29. G. Zhong, L. Liu, T. Fan, P. Fan, H. Ji, Degradation of transcription factor RFX5 during the inhibition of both constitutive and interferon gamma-inducible major histocompatibility complex class I expression in chlamydia-infected cells. The Journal of experimental medicine 191, 1525–1534 (2000).

30. G. Zhong, Killing me softly: chlamydial use of proteolysis for evading host defenses. Trends in microbiology 17, 467–474 (2009).

31. G. Zhong, Chlamydia trachomatis secretion of proteases for manipulating host signaling pathways. Frontiers in microbiology 2, 14 (2011).

32. Z. Yang, L. Tang, X. Sun, J. Chai, G. Zhong, Characterization of CPAF critical residues and secretion during Chlamydia trachomatis infection. Infection and immunity 83, 2234–2241 (2015).

33. H. Feng et al., Transferrin Receptor Is a Specific Ferroptosis Marker. Cell reports 30, 3411–3423.e3417 (2020).

34. W. Wang et al., CD8(+) T cells regulate tumour ferroptosis during cancer immunotherapy. Nature 569, 270–274 (2019).

35. X. Fang et al., Ferroptosis as a target for protection against cardiomyopathy. Proc Natl Acad Sci U S A 116, 2672–2680 (2019).

36. Q. Hu et al., GPX4 and vitamin E cooperatively protect hematopoietic stem and progenitor cells from lipid peroxidation and ferroptosis. Cell death & disease 12, 706 (2021).

37. M. Matsushita et al., T cell lipid peroxidation induces ferroptosis and prevents immunity to infection. The Journal of experimental medicine 212, 555–568 (2015).

38. A. L. Chen, K. A. Johnson, J. K. Lee, C. Sütterlin, M. Tan, CPAF: a Chlamydial protease in search of an authentic substrate. PLoS pathogens 8, e1002842 (2012).

39. S. A. Paschen et al., Cytopathicity of Chlamydia is largely reproduced by expression of a single chlamydial protease. The Journal of cell biology 182, 117–127 (2008).

40. A. E. Carlisle et al., Selenium detoxification is required for cancer-cell survival. Nature metabolism 2, 603–611 (2020).

